# CaMKKβ regulates transcription factor Elf2 gene methylation to maintain endothelial junctional barrier integrity

**DOI:** 10.1101/2025.02.27.640631

**Authors:** Dong-Mei Wang, Juan Yang, Mohammad Owais Ansari, Zarema Arbieva, Mark Maienschein-Cline, Viswanathan Natarajan, Asrar B. Malik, Chinnaswamy Tiruppathi

## Abstract

The expression of endothelial-enriched receptor tyrosine kinase Tie2 and vascular endothelial cadherin VE-cadherin in endothelial cells (EC) is essential for maintaining endothelial barrier integrity and vascular homeostasis. The Ets family of transcription factors plays a critical role in regulating the expression of Tie2 and VE-cadherin in EC. However, the transcriptional regulation of these factors remains poorly understood. In this study, we performed whole genome methylation analysis and found that the gene encoding the Ets family transcription factor Elf2/Nerf2 was hypermethylated in Camkkβ-deficient (*Camkkβ^−/−^*) mice. Notably, these mice exhibited significantly reduced mRNA and protein levels of Elf2, Tie2, and VE-cadherin, along with uncontrolled lung vascular injury following LPS challenge. Additionally, either depletion of CaMKKβ in human lung microvascular endothelial cells or EC-specific deletion of *Camkkβ* in mice led to a decrease in the expression of Elf2, Tie2, and VE-cadherin. Furthermore, administration of the DNA methyltransferase inhibitor 5-azacytidine (5-AZA) or EC-specific expression of wild-type CaMKKβ but not the kinase-defective mutant restored the expression of Elf2, Tie2, and VE-cadherin in *Camkkβ^−/−^*mice. We find that under baseline conditions, methyl CpG binding protein MeCP2 repressed Elf2 expression in EC. Importantly, in EC-restricted *Elf2* knockout mice (*Elf2^1EC^*), lung endothelial barrier integrity was compromised due to dramatically reduced levels of Tie2 and VE-cadherin. Together, these findings underscore the crucial role of CaMKKβ in regulating endothelial junctional barrier integrity by controlling Elf2 expression.

## Introduction

Vascular endothelial cadherin (VE-cadherin), expressed at endothelial adherens junctions (AJs), is essential for maintaining vascular integrity and endothelial homeostasis (1–5). VE-cadherin forms Ca²⁺-dependent homophilic cis and trans dimers at AJs, establishing tight cell-to-cell adhesions that prevent the free passage of plasma and blood constituents. The receptor tyrosine kinase Tie2 is highly expressed in endothelial cells (EC) and is crucial for angiogenesis and vascular barrier maintenance (6). Tie2 binds the ligands Angiopoietin-1 (Ang-1) and Angiopoietin-2 (Ang-2), producing opposing effects on vascular barrier stability (6–8). Ang-1, produced by pericytes and platelets, prevents vascular leak by interacting with Tie2, whereas Ang-2, primarily released from EC during sepsis, promotes vascular leak by antagonizing Tie2 function (6–8). The Ang-1/Tie2 signaling axis is constitutively active in quiescent blood vessels, stabilizing the endothelial barrier by increasing VE-cadherin expression at AJs through Rac1-mediated inhibition of RhoA (6,9,10). Activation of Tie2 with recombinant Ang-1 has been shown to block sepsis-induced vascular leak (11,12), highlighting the importance of Tie2 signaling in stabilizing the endothelial junctional barrier. However, the expression of both VE-cadherin and Tie2 in EC is downregulated in inflammatory conditions, e.g., sepsis (13–15), underscoring the need to investigate transcriptional mechanisms that regulate these critical endothelial barrier proteins.

The Ets family of transcription factors, which bind the DNA sequence 5’-GGA(A/T)-3’ via their conserved ETS domain, plays a key role in the expression of endothelial-specific genes (16,17). Nineteen Ets factors have been identified in EC, many of which are essential for vascular development and angiogenesis (17–19). Dube et al. (20) demonstrated that the Ets transcription factor Elf2 transactivates the Tie2 promoter, while other Ets factors had no effect on Tie2 transcription. Additionally, evidence suggests that Elf2 promotes VE-cadherin transcription (21); however, the role of EC-expressed Elf2 in regulating the expression of Tie2 and VE-cadherin is not well understood.

CaMKKβ is a Ser/Thr kinase activated by the Ca²⁺-signal transducer calmodulin (CaM) (22,23). It activates several downstream kinases, including CaM-kinase I (CaMKI), CaM-kinase IV (CaMKIV), protein kinase B (PKB/Akt), and 5’AMP-activated protein kinase (AMPK), through phosphorylation in response to increased intracellular Ca²⁺ levels (22,23). CaMKKβ is also involved in transcriptional activation by phosphorylating transcription factors and gene repressors (23–26). It has been shown that CaMKKβ-mediated phosphorylation of SIRT1 in endothelium prevents atherosclerosis in mice (27). Further, recent evidence suggests that under ischemic conditions, CaMKKβ protects EC and the blood-brain barrier through SIRT1 phosphorylation and activation (28). Given that CaMKKβ inhibits endothelial dysfunction and promotes gene transcription, we hypothesized that EC-expressed CaMKKβ regulates the endothelial junctional barrier by controlling the expression of barrier-stabilizing genes. We found that the gene encoding the Ets transcription factor Elf2 was hypermethylated in CaMKKβ deficient (*Camkkβ^−/−^*) mice. Consistent with this, the expression of Elf2, Tie2, and VE-cadherin at both mRNA and protein levels was significantly reduced in *Camkkβ^−/−^*mice, which also exhibited uncontrolled lung vascular injury following low-dose LPS challenge. Additionally, either depletion of CaMKKβ in human lung microvascular endothelial cells (HLMVEC) or EC-specific deletion of CaMKKβ in mice suppressed Elf2, Tie2, and VE-cadherin expression. Further in EC-restricted Elf2 deletion (*Elf2^ΔEC^*) in adult mice, we observed that the expression of Tie2 and VE-cadherin was suppressed, and the endothelial junctional barrier was compromised. These findings underscore the critical role of CaMKKβ-mediated Elf2 expression in regulating endothelial junctional barrier through the expression of Tie2 and VE-cadherin in EC.

## Results

### CaMKKβ deficiency promotes uncontrolled lung vascular inflammation in mice

*Camkkβ^+/−^* (heterozygous) mice were bred to generate homozygous (*Camkkβ^−/−^*) mice. Immunoblot (IB) analysis of lung tissue confirmed the absence of CaMKKβ expression in *Camkkβ^−/−^*mice (**Fig. 1a**). Male and female *Camkkβ^−/−^* and wildtype (*Camkkβ^+/+^*) mice aged 8 to 12 weeks were used for experiments. A low-dose LPS (5 mg/kg, i.p.) challenge resulted in 100% mortality in *Camkkβ^−/−^* mice within 48 hours, while WT mice showed no mortality (**Fig. 1b**). Polymicrobial sepsis induced by cecal ligation and puncture (CLP) also caused 100% mortality in *Camkkβ^−/−^* mice within 24 hours (**Fig. 1c**). Hematoxylin-and-eosin (H&E) staining of lung sections (**Fig. 1d**) and myeloperoxidase (MPO) activity measurements (**Fig. 1e**) demonstrated enhanced lung vascular injury in *Camkkβ^−/−^* mice. We also observed a significantly greater vascular leak in response to LPS and PAR-1 activating peptide in *Camkkβ^−/−^* mice compared to WT mice (**Fig. 1f, g**). These results suggest a potential link between CaMKKβ deficiency and endothelial barrier dysfunction.

**Fig. 1:**
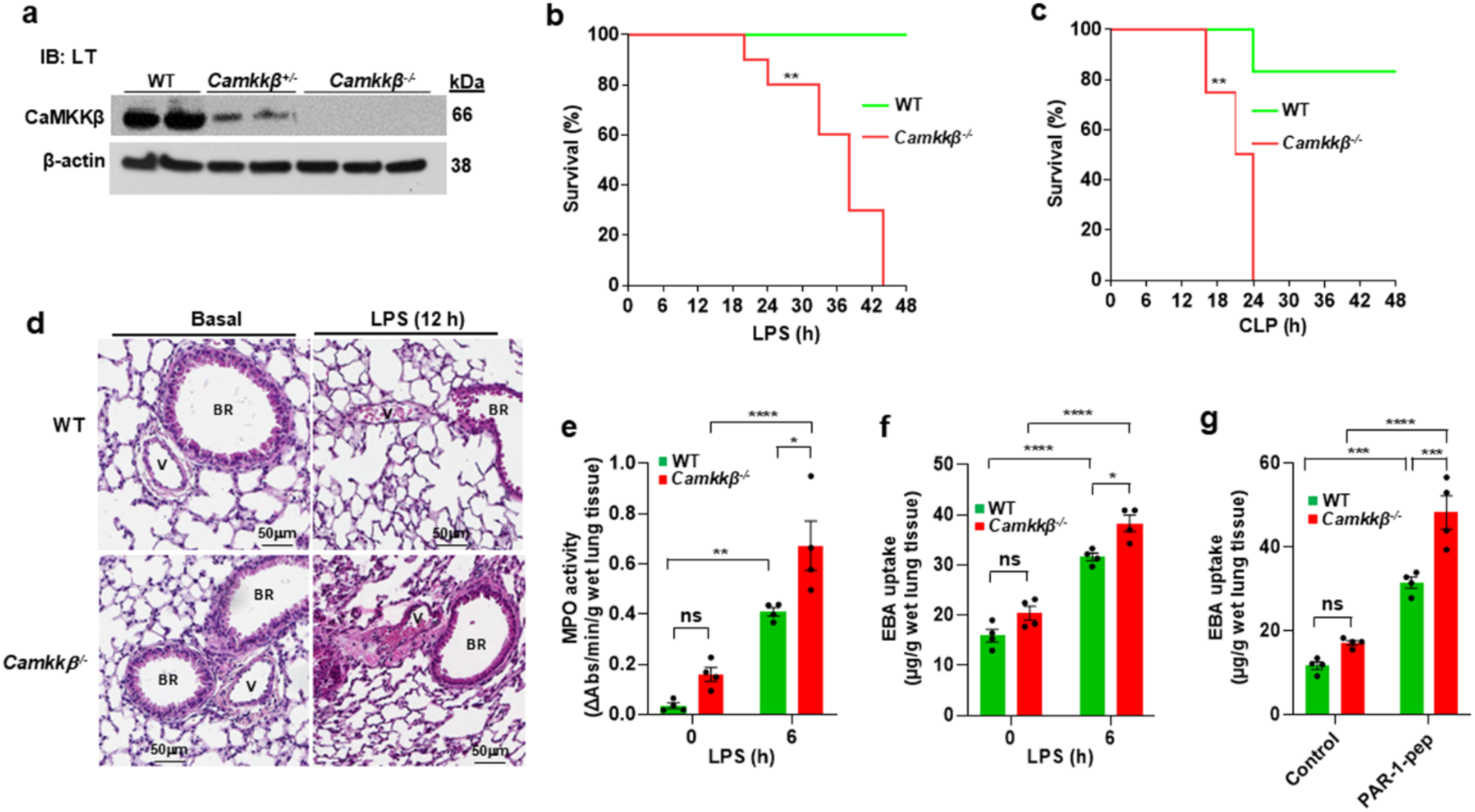
CaMKKβ deficiency promotes uncontrolled lung vascular inflammation in mice. **a**, Immunoblot (IB) showing CaMKKβ expression in lung tissue (LT) from WT (*Camkkβ^+/+^*), *Camkkβ^+/−^*, and *Camkkβ^−/−^* mice. **b**, Survival following LPS challenge (5 mg/kg; i.p.) in WT and *Camkkβ^−/−^* mice. N = 10 in each group. **p<0.01 (log-rank test). **c**, Survival following CLP in WT and *Camkkβ^−/−^* mice. N = 8 in each group. **p<0.01 (log-rank test). **d**, H&E staining of lung sections from WT and *Camkkβ^−/−^* mice challenged with LPS (5 mg/kg; i.p.). Br, bronchi; V, blood vessel. **e**, After LPS challenge (5 mg/kg, i.p), lungs harvested were used to measure MPO activity. **f**, 0 h and 6 h post LPS challenge, *in vivo* lung vascular leak was assessed by Evans blue bound albumin (EBA) uptake. **g**, PAR-1-activating peptide (TFLLRNPNDK-NH2)-induced *in vivo* lung vascular leak was measured for a period of 1 h in WT and *Camkkβ^−/−^* mice. **e-g**, N= 4 per genotype per time point; ns: not significant, *p<0.05, **p<0.01, ***p< 0.001, ****p< 0.0001, WT vs *Camkkβ^−/−^* (Two-way ANOVA).

### CaMKKβ regulates genes critical for endothelial barrier integrity via DNA methylation

Given that VE-cadherin and Tie2 are essential for endothelial barrier integrity (1,14), we assessed their expression. Protein levels of VE-cadherin and Tie2 were significantly reduced in lung tissues of *Camkkβ^−/−^* mice compared to *Camkkβ^+/+^*and *Camkkβ^+/−^* mice (**Fig. 2a**). The mRNA levels of VE-cadherin, Tie2, and the Tie2 agonist Ang-1 were also decreased in *Camkkβ^−/−^*mice, while mRNA of the Tie2 antagonist Ang-2 was increased (**Fig. 2b**). We also observed that the expression of mRNA for β-catenin, VE-cadherin associated junctional protein, was reduced; however, mRNA levels for Occludin, VEGFR1, and VEGFR2 were not significantly altered (**Fig. 2b**). These findings imply that CaMKKβ is required for the expression of VE-cadherin and Tie2 in EC. Next to study whether kinase function of CaMKKβ is required for the expression of VE-cadherin and Tie2, we i.v. injected *Camkkβ^−/−^* mice with liposome-CaMKKβ plasmid complexes containing either WT-CaMKKβ or a kinase-defective CaMKKβ mutant (K193A-CaMKKβ). We observed only that WT-CaMKKβ rescued the expression of VE-cadherin and Tie2 (**Fig. 2c**). This underscores the role of CaMKKβ’s kinase activity in regulating the transcription of these genes.

**Fig. 2:**
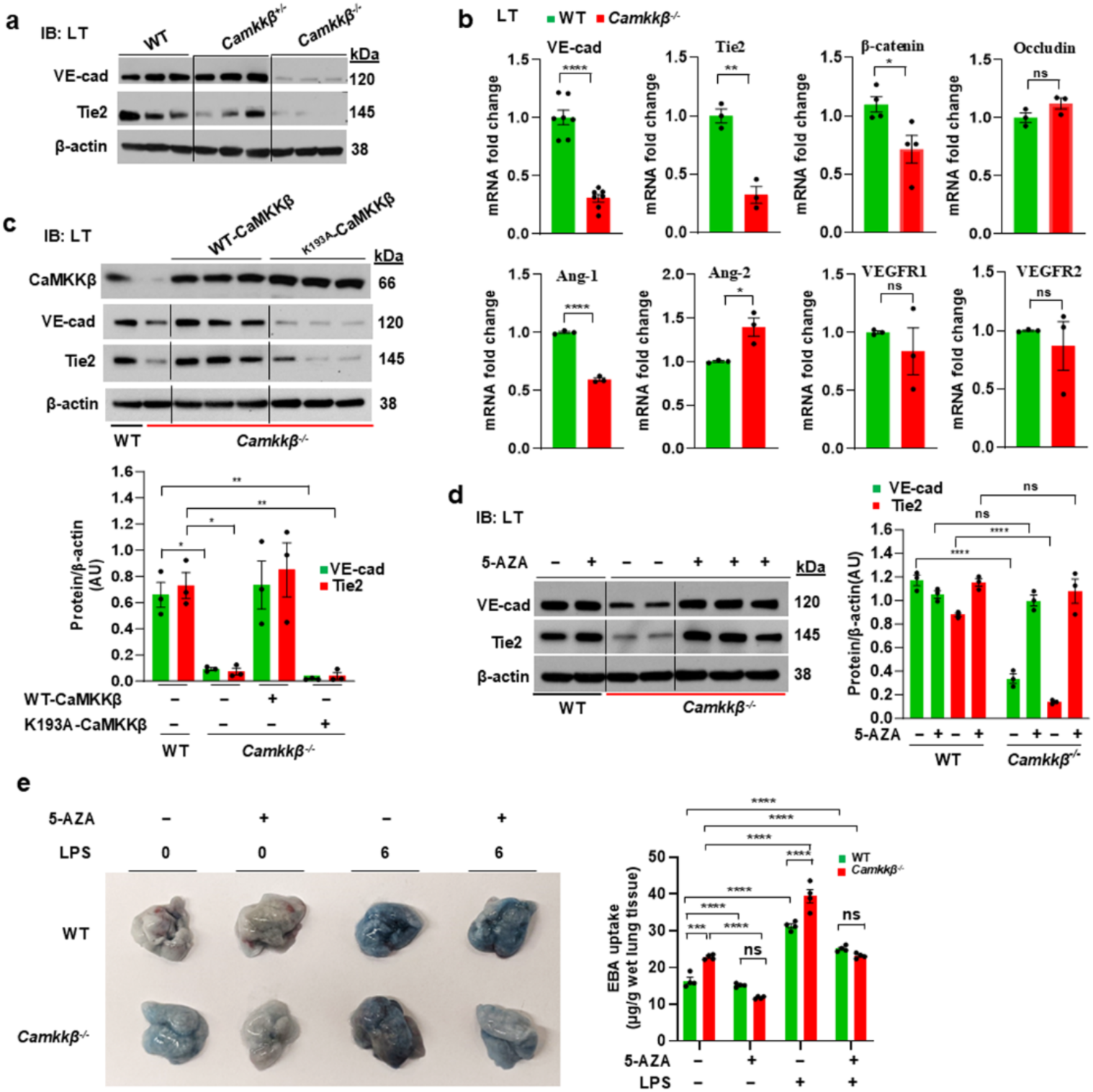
CaMKKβ regulates genes critical for endothelial integrity via DNA methylation. **a,** Defective expression of VE-cadherin and Tie2 in *Camkkβ^−/−^* mice. Total lung lysates from WT, *Camkkβ^+/−^,* and *Camkkβ^−/−^* mice were used for IB. Representative blot is shown; N = 3 mice/genotype. **b**, CaMKKβ regulates transcription of genes responsible for endothelial barrier integrity. Lung tissue (LT) mRNA levels were determined by qRT-PCR. N ζ 3 mice per group; ns, not significant; *p< 0.05, **p< 0.01, ****p< 0.0001, WT vs *Camkkβ^−/−^* (Unpaired *t* test). **c**, Expression of WT-CaMKKβ in lung endothelial cells of *Camkkβ^−/−^* mice rescued the expression of VE-cadherin and Tie2. *Camkkβ^−/−^* mice were injected with liposome-CaMKKβ plasmid (1 µg plasmid/gm body weight) complexes containing either WT-CaMKKβ or the kinase defective CaMKKβ (^K193A^-CaMKKβ) mutant. At 72 h after injection, lungs harvested were used for IB analysis. N = 3 mice per group; representative blot is shown. **d**, DNA methyltransferase inhibition restored expression of VE-cadherin and Tie2 in *Camkkβ^−/−^* mice. WT and *Camkkβ^−/−^* mice were injected with 5-AZA (1 mg/kg; i.p.) or vehicle for 5 consecutive days and lungs harvested used for IB analysis. N = 3 mice per group; representative blot is shown; n.s, not significant; ****p< 0.0001. *p< 0.05, **p< 0.01, WT vs WT-CaMKKβ or ^K193A^-CaMKKβ. **e**, WT and *Camkkβ^−/−^* mice were injected with 5-AZA as above were treated with LPS (5 mg/kg, i.p.) for 0 and 6 h and lung vascular permeability was determined via measuring EBA uptake. N= 4 mice/group; n.s, not significant, ***p< 0.001, ****p< 0.0001.

To investigate the underlying mechanism involved in the transcription of VE-cadherin and Tie2, we explored DNA methylation. Treatment with the DNA methyltransferase inhibitor 5-azacytidine (5-AZA) for 5 days restored VE-cadherin and Tie2 expression in *Camkkβ^−/−^* mice (**Fig. 2d**). We also measured LPS-induced lung vascular permeability after 5-AZA treatment. We noted that 5-AZA treatment reduced LPS-induced lung vascular permeability in *Camkkβ^−/−^* mice (**Fig. 2e**). These findings support the notion that DNA methylation represses VE-cadherin and Tie2 transcription in the absence of CaMKKβ.

### CaMKKβ deficiency leads to hyper-methylation of the Elf2 transcription factor gene

Given the involvement of CaMKKβ in the regulation of VE-cadherin and Tie2, we examined the DNA methylation status of these genes. Our analysis revealed low levels of methylation-prone CpG sites in the promoter regions of VE-cadherin and Tie2 (**Fig. 3a**). Genome-wide methylation analysis using MeDIP-seq identified aberrant methylation of several genes in *Camkkβ^−/−^* mice, including two highly methylated genes, Pbk and Elf2 (**Fig. 3b-c; Supplemental data Table 1**). Given that PBK/TOPK’s role in endothelial function is poorly understood, we focused on the transcription factor Elf2, which is known to activate transcription of Tie2 and VE-cadherin in EC. Consistent with methylation data, both mouse and human Elf2 genes were enriched with CpG sequences (**Fig. 3d**). The mRNA and protein expression of Elf2, but not Elf1, were suppressed in *Camkkβ^−/−^* mice (**Fig. 3e, f**). Treatment with 5-AZA or expression of WT-CaMKKβ (but not kinase-defective CaMKKβ) restored Elf2 expression in *Camkkβ^−/−^* mice (**Fig. 3g**), suggesting that DNA methylation mediates Elf2 repression in the absence of CaMKKβ.

**Fig. 3:**
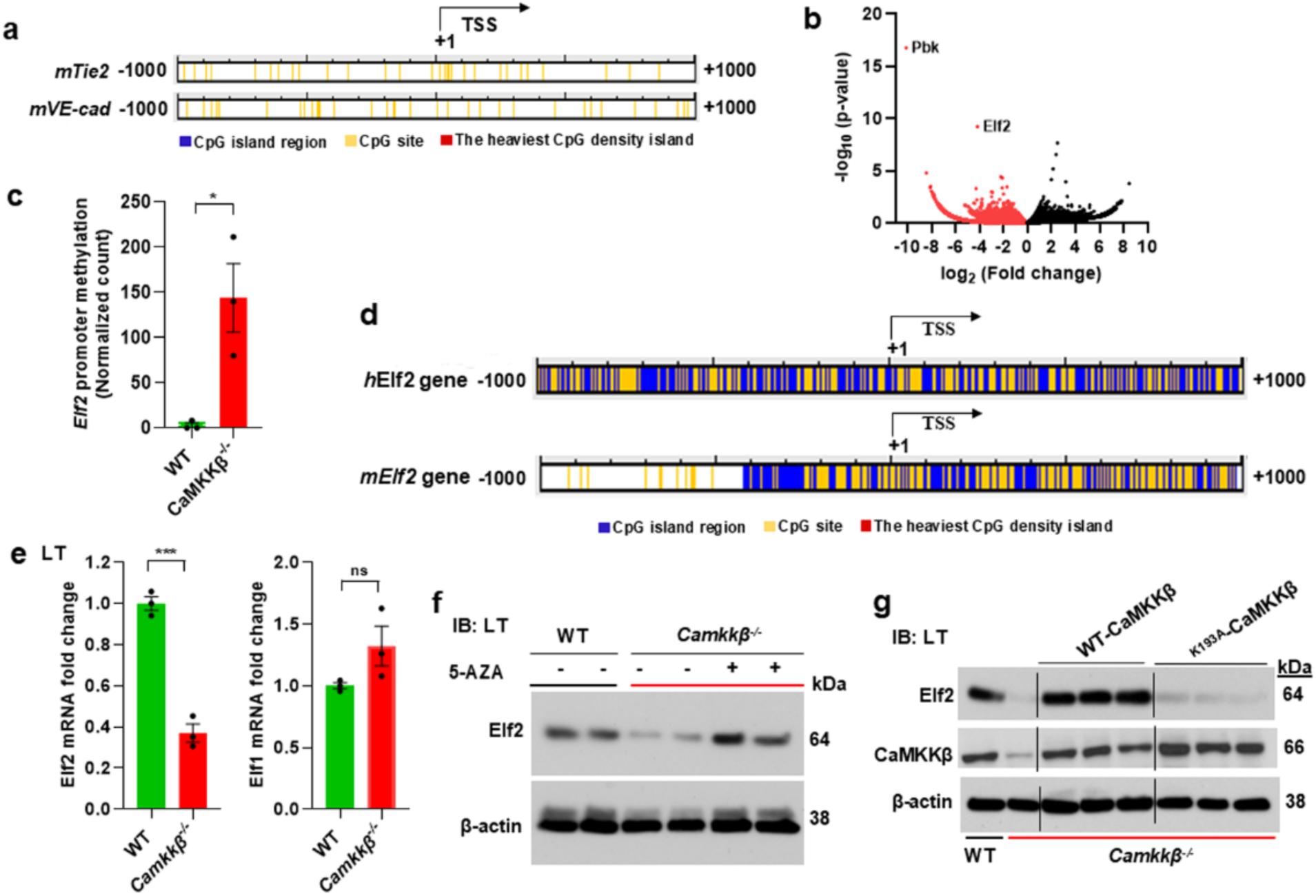
CaMKKβ deficiency leads to hyper-methylation of the Elf2 transcription factor gene. **a,** DNA sequence reveals the low levels of CpG sites in the promoter regions of mouse Tie2 and VE-cadherin (Cdh5). **b**, Volcano plot showing changes in methylation level of genes in *Camkkβ^+/+^* and *CamKKβ^−/−^* mice. The ratio of CpG methylated sites in genes (+/– 2kb of the TSS) of *Camkkβ^+/+^/Camkkβ^−/−^* mice were presented (fold changes *vs*. p-value). **c**, Quantified data showing hyper-methylation of Elf2 gene promoter in *CamKKβ^−/−^* mice. N = 3 samples per genotype; *p<0.05 (unpaired *t* test). **d**, DNA sequence analysis reveals the abundance of CpG islands and CpG sites in the promoter regions of human and mouse Elf2 genes. **e**, WT and *Camkkβ^−/−^* mice lung mRNA levels of Elf2 and Elf1 were determined by qRT-PCR. N = 3 mice per genotype; ns, not significant, ***p<0.001 (unpaired *t* test). **f**, DNA methyl transferase inhibition restores Elf2 expression in *Camkkβ^−/−^* mice. WT and *Camkkβ^−/−^* mice were injected with 5-AZA (1 mg/kg, i.p.) or vehicle for 5 days as described above in Fig. 2c. Lungs were harvested and used for IB. N = 3 mice per group; representative blot is shown. **g**, Expression of WT-CaMKKβ in EC of *Camkkβ^−/−^* mice rescued Elf2 expression. *Camkkβ^−/−^* mice were injected with liposome-CaMKKβ plasmid (1 µg plasmid/g body weight) complexes containing either WT-CaMKKβ or the kinase defective CaMKKβ (^K193A^-CaMKKβ) mutant. At 72 h after injection, lungs were harvested and used for IB analysis. N = 3 mice per group; representative blot is shown.

### CaMKKβ regulates Elf2, Tie2, and VE-cadherin expression in EC

To determine the role of EC-specific CaMKKβ in Elf2 regulation, we disrupted CaMKKβ in mouse lung vascular endothelial cells using a CRISPR/Cas9 system. This resulted in reduced expression of Tie2, VE-cadherin, and Elf2 in the lungs of WT mice (**Fig. 4a**). Similarly, suppression of CaMKKβ expression in HLMVEC using siRNA led to a marked reduction in the expression of these genes (**Fig. 4b**). Immunostaining showed that LPS challenge disrupted VE-cadherin at cell-cell junctions, with its expression restored at 24 hours in control HLMVEC but not in CaMKKβ-depleted HLMVEC (**Fig. 4c**). These data underscore CaMKKβ’s critical role in maintaining endothelial barrier integrity. We further investigated the role of CaMKKβ signaling during vascular injury. Following LPS challenge (10 mg/kg, i.p.) of WT mice, we observed that the expression of CaMKKβ, Elf2, Tie2, and VE-cadherin was significantly downregulated at 6-12 hours but restored to baseline by 48 hours (**Fig. 4d, e**). This highlights the role of CaMKKβ in signaling for the re-expression of these genes post-vascular injury.

**Fig. 4.**
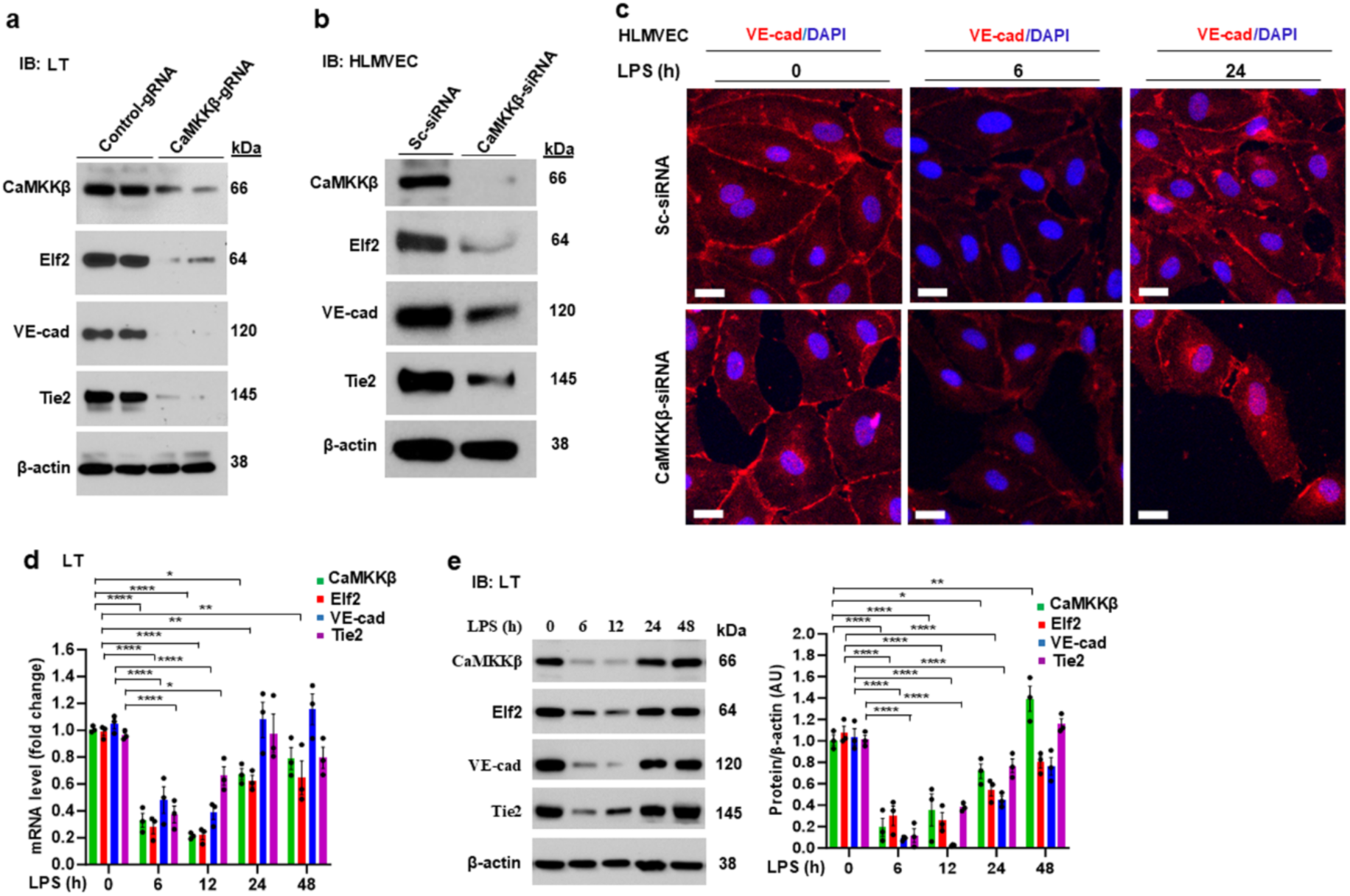
CaMKKβ regulates Elf2, VE-cadherin, and Tie2 expression in EC. **a**, EC-specific disruption of *Camkkβ* in mice inhibits expression of Elf2, Tie2, and VE-cadherin. WT mice were injected with CRISPR plasmid encoding control-sgRNA (5′-GCGAGGTATTCGGCTCCGCG-3′) or *m*CaMKKβ-sgRNA (5′-AAACGTCGGATCAGCTTCTTTTTGC-3′). The liposomes and CRISPR plasmid mixture was injected i.v. into the mouse (1 µg plasmid/g body weight). Lungs harvested at 96 h after injection were used for IB analysis. Representative blots are shown (N = 4 mice/group). **b-c**, HLMVEC were transfected with 100 nM of either scrambled-siRNA (Sc-siRNA) or CaMKKβ-siRNA. At 72 h after transfection, cell lysates were used for IB analysis (**b**) or challenged with LPS (5 μg/ml) for the indicated times and then cells were stained with anti-VE-cadherin pAb and DAPI (scale bar = 20 μm) (**c**). **b**, Results show a representative blot. **d-e**, TLR4 signaling regulates endothelial expression of Tie2 and VE-cadherin through CaMKKβ-mediated Elf2 transcription. WT mice were challenged with LPS (10 mg/kg, i.p.) for different time intervals. After LPS challenge, lungs harvested were used for qRT-PCR (**d**) and IB analysis (**e**). **e**, results show a representative blot. **d** and **e**, Shown are mean values ± SEM (n =3 mice/time point). *p<0.05; **p<0.01; ****p<0.0001 (Two-way ANOVA).

### CaMKKβ inhibits MeCP2-mediated repression of Elf2 transcription in EC

Methyl CpG binding protein MeCP2, known to induce long-term silencing of genes in neurons (25,29) is expressed in EC (30,31). MeCP2 has conserved S^80^ in human, rat, and mouse, and this S^80^ is phosphorylated under basal state, which promotes MeCP2 binding to methylated CpG DNA (29,32). Ca^2+^ influx causes phosphorylation of MeCP2 at S^421^ to dissociate MeCP2 from methylated CpG DNA and hence promotes gene transcription (25,26), but it is not known whether it has any role in regulating transcription of Elf2 in EC. Thus, we investigated whether Ca^2+^/CaM-dependent kinase CaMKKβ is required for the expression of *Elf2* through inhibiting MeCP2 binding to methylated CpG sites. In HLMVEC, we observed that under baseline conditions, MeCP2 is mainly localized in the nucleus (**Fig. 2a, b**). Further, we observed thrombin or LPS challenge caused dephosphorylation of S^80^ and phosphorylation of S^421^ in MeCP2 in HLMVEC (**Fig. 5c, d).** In support of these findings, inhibition of CaMKKβ, prevented thrombin or LPS induced phosphorylation of MeCP2 at S^421^ (**Fig. 5c, d**) indicating that Ca^2+^ influx induced CaMKKβ activation caused phosphorylation of MeCP2 at S^421^ in EC. Next to address the role of nuclear localized MeCP2, we determined MeCP2 binding to Elf2 promoter by chromatin immunoprecipitation (ChIP) assay. Here, we observed the association of MeCP2 with the upstream proximal promoter regions (─307 and ─561) of *h*Elf2 gene in HLMVEC under basal state (**Fig. 5e**). Interestingly, thrombin stimulation, which induces Ca^2+^ influx in EC, caused dissociation of MeCP2 from the *h*Elf2 promoter (**Fig. 5e**) indicating the potential role of MeCP2 in suppressing the transcription of Elf2.

**Fig. 5:**
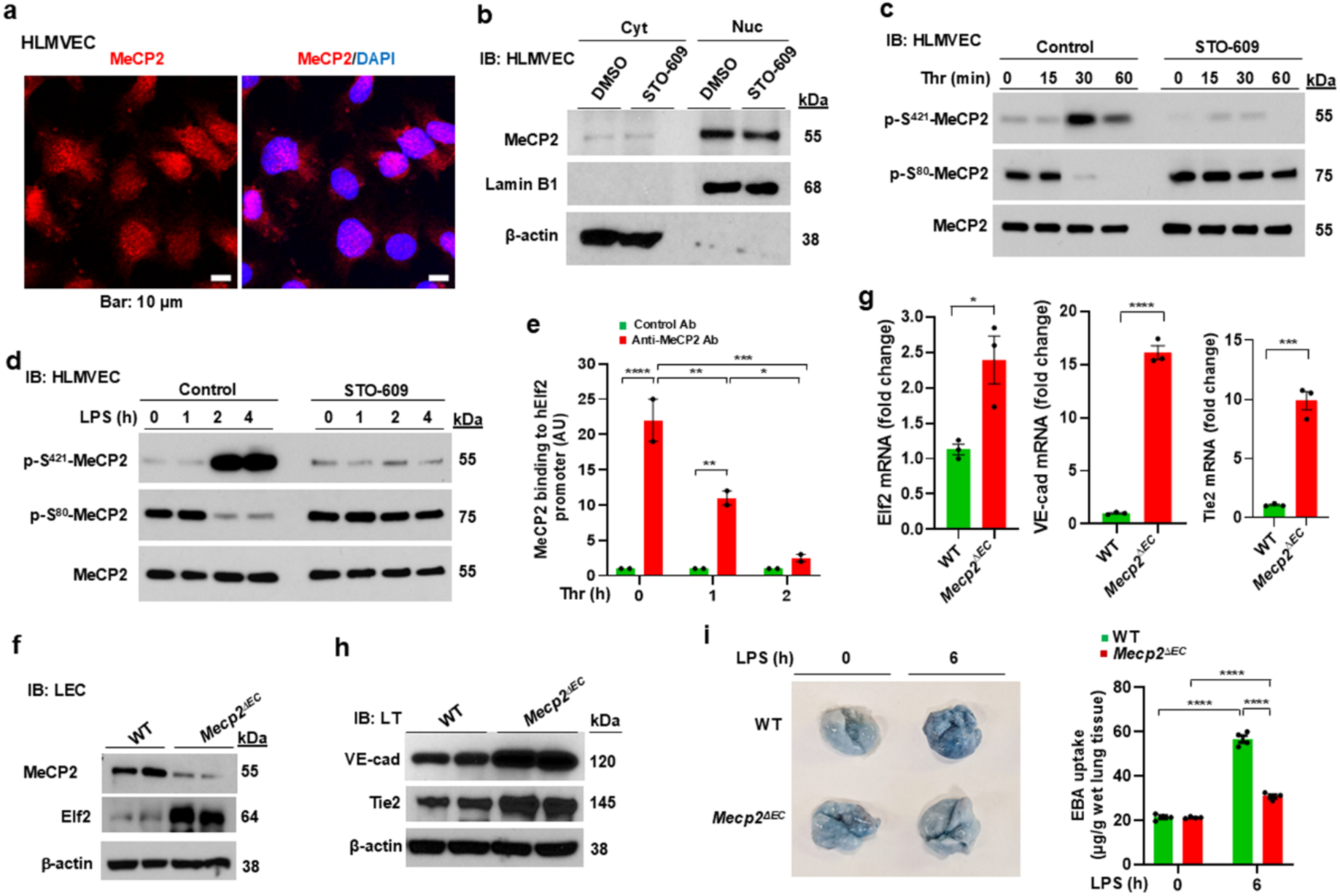
a-e, CaMKKβ inhibits MeCP2-mediated repression of Elf2 transcription in EC. **a**, HLMVEC incubated with serum free medium for 12 h were stained with anti-MeCP2 pAb and DAPI. **b**, HLMVEC incubated with serum free medium for 12 h and then incubated with and without CaMKKβ inhibitor STO-609 (1 μM) for 60 min. Thereafter cytosolic and nuclear fractions prepared were used for IB. **c-d**, HLMVEC pretreated with STO-609 (1 μM) or vehicle (DMSO) for 60 min were exposed to thrombin (25 nM) or LPS (5 μg/ml) for different time intervals and used for IB analysis to determine phosphorylation of MeCP2 at S421 and S80. Results shown are representative blots of 2 independent experiments. **e**, MeCP2 was associated with the Elf2 gene promoter under basal state in HLMVEC. HLMVEC grown to confluency were used for ChIP assay to determine MeCP2 interaction with the Elf2 promoter. Thrombin stimulation caused the dissociation of MeCP2 from the promoter regions of Elf2 gene. Results shown are mean values ± SEM of two experiments. *p<0.05; **p<0.01; ***p<0.001; ****p<0.0001 (Two-way ANOVA). **f-I**, **EC-restricted Mecp2 deletion (*Mecp2^1EC^*) in mice augments expression of Elf2 and mitigated LPS-induced lung vascular leak. f**, Lung endothelial cells (LEC) isolated from WT and *Mecp2^1EC^* mice were used for IB analysis to determine the expression of MeCP2 and Elf2. Results showed the loss of MeCP2 and augmented expression of Elf2 in *Mecp2^1EC^* mice LEC. **g**, Lungs harvested from WT and *Mecp2^1EC^*mice were used for qRT-PCR to determine expression of mRNA for Elf2, VE-cadherin and Tie2. N = 3 mice/genotype; *p<0.05, ***p<0.001; ****p<0.0001 (unpaired *t* test). **h,** Lungs were used for IB to determine the expression of VE-cadherin and Tie2. **i**, WT and *Mecp2^1EC^*mice were challenged with LPS (10 mg/kg; i.p.) and then used to assess *in vivo* lung vascular leak by measuring EBA uptake. N = 4 mice per genotype; **p<0.01; ****p< 0.0001 (Two-way ANOVA).

To study the *in vivo* role of EC-expressed MeCP2 in repressing Elf2 transcription, we generated an EC-restricted *Mecp2* knockout (*Mecp2^1EC^*) mice by crossing loxP-flanked *Mecp2* with Cdh5-Cre (VE-cadherin-Cre) mice. Lung endothelial cells (LEC) isolated from *Mecp2^fl/fl^* (WT) and *Mecp2^1EC^* mice showed the dramatically reduced expression of MeCP2 and increased expression of Elf2 in *Mecp2^1EC^* mice (**Fig. 5f**). Importantly, we observed that baseline expression of mRNA for Elf2, Tie2, and VE-cadherin was significantly increased in *Mecp2^1EC^* mice compared to WT mice (**Fig. 5g**). Also, we noted that expression of protein for Tie2 and VE-cadherin was increased in *Mecp2^1EC^* mice (**Fig. 5h**). In support of these results, LPS-induced lung vascular permeability was markedly reduced in *Mecp2^1EC^* mice compared with wildtype counterparts (**Fig. 5i).** These results support the proposal that EC-expressed MeCP2 plays a crucial role in repressing Elf2 expression.

### Elf2-mediated expression of Tie2 and VE-cadherin contributes to endothelial barrier integrity in human lung EC

To establish a link between Elf2 expression and the transcription of Tie2 and VE-cadherin, we analyzed the 5′-regulatory regions of the mouse and human genes encoding these proteins. Multiple potential Elf2 (Ets) binding sites were identified in the promoter regions of both human and mouse genes (**Fig. 6a; mouse data not shown**). Using a ChIP assay, we identified Elf2 binding to sites in the Tie2 and Cdh5 (VE-cadherin) promoter regions under baseline conditions in HLMVEC (**Fig. 6b, c**). Following LPS challenge, Elf2 was dissociated from these sites in these promoter regions, with the maximal dissociation observed at 6 hours post-challenge. Elf2 binding was restored to near basal level within 24 hours (**Fig. 6b, c**), consistent with the LPS-induced downregulation of Elf2 observed in WT mice (**Fig. 4d, e**).

**Figure 6.**
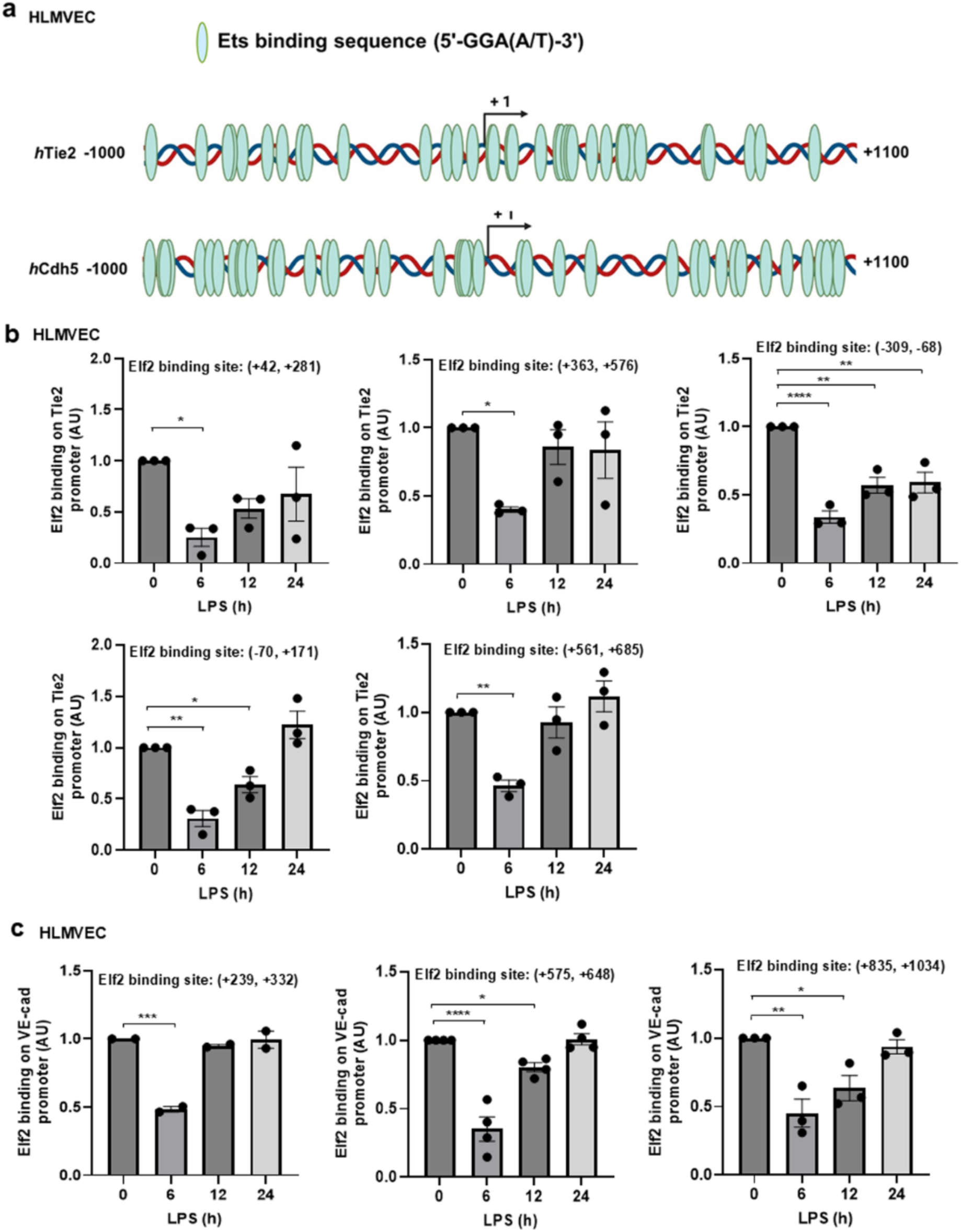
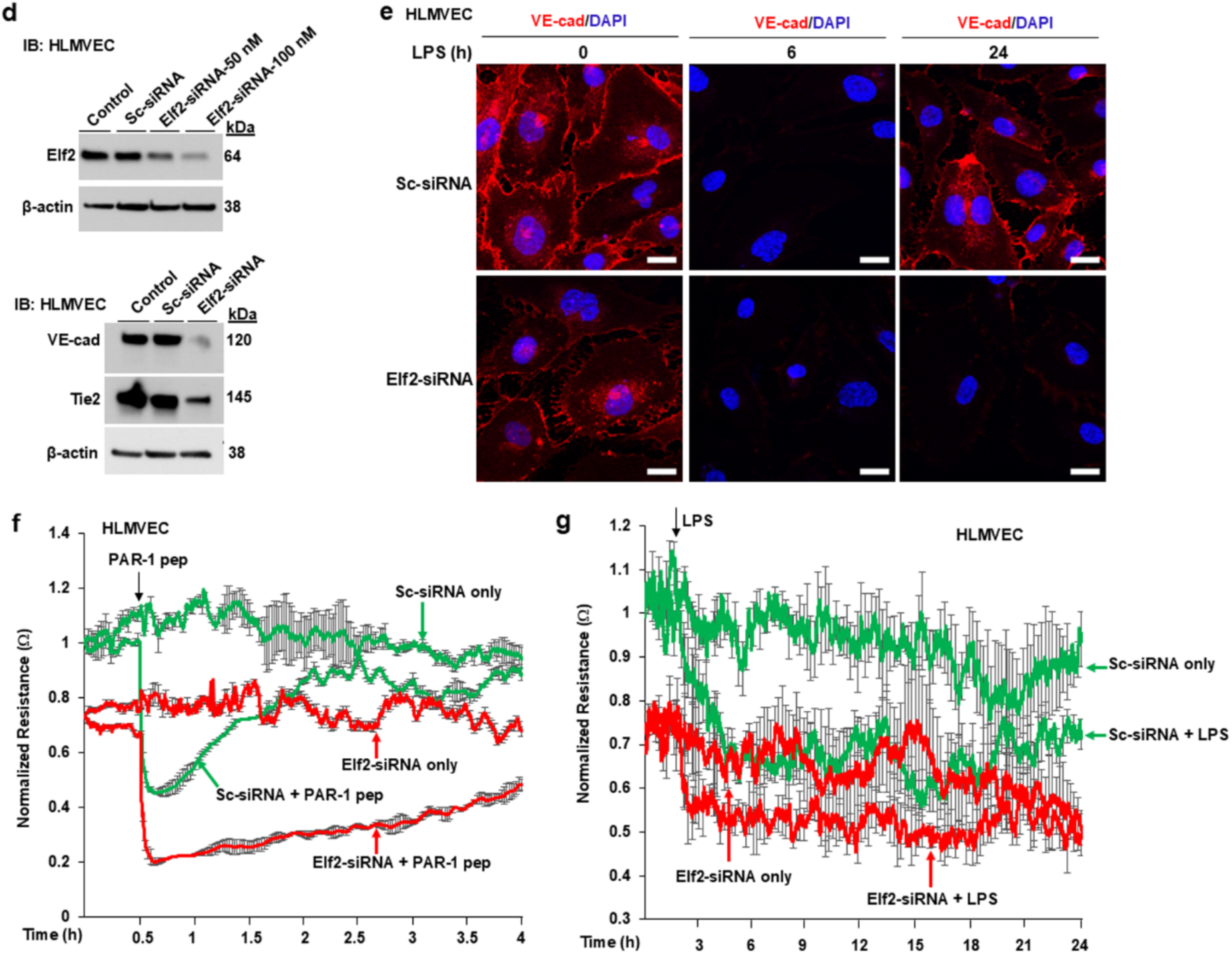
Elf2-mediated expression of Tie2 and VE-cadherin contributes to endothelial barrier integrity in human lung EC. **a**, Schematics showing Ets binding sites in the promoter regions of *h*Tie2 and *h*Cdh5. **b-c**, HLMVEC challenged with LPS (5 μg/ml) for different time intervals were used for ChIP assay to determine Elf2 binding to the promoter regions of Tie2 (**b**) and of VE-cadherin (Cdh5) (**c**). Results shown are mean values of three experiments. *p<0.05; **p<0.01; ***p<0.001; ****p<0.0001 (One-way ANOVA). **d**, HLMVEC transfected with scrambled-siRNA (Sc-siRNA) or *h*Elf2-siRNA were used for IB analysis. **e**, HLMVEC transfected with Sc-siRNA or *h*Elf2-siRNA were stained with anti-VE-cadherin Ab and DAPI to assess endothelial AJ integrity (scale bar = 20 μm). **f** and **g**, HLMVEC transfected with Sc-siRNA or *h*Elf2-siRNA were used to measure PAR-1 activating peptide (25 μM)- or LPS (5 µg/ml)-induced changes in TER as an assessment of endothelial permeability. Arrow indicates time at which PAR-1 activating peptide (PAR-1 pep), LPS or media was added.

To further explore the role of Elf2, we silenced its expression using siRNA specific to Elf2 (Elf2-siRNA) in HLMVEC. Elf2 silencing significantly reduced the expression of both VE-cadherin and Tie2 compared to scrambled siRNA (Sc-siRNA)-transfected cells (**Fig. 6d**). Immunostaining revealed decreased VE-cadherin expression at AJs under baseline conditions in Elf2-siRNA-treated cells compared to controls (**Fig. 6e**). After LPS challenge, VE-cadherin was lost from AJs in both Elf2-siRNA and Sc-siRNA-transfected cells at 6 hours. The VE-cadherin expression was restored in Sc-siRNA-treated cells by 24 hours but remained suppressed in Elf2-siRNA-treated cells, with additional EC loss observed (**Fig. 6e**). Endothelial permeability, measured by transendothelial monolayer electrical resistance (TER), was significantly lower in Elf2-siRNA-transfected HLMVEC under baseline conditions (**Fig. 6f, g**). Both PAR-1 activating peptide and LPS caused a more pronounced reduction in TER over time in Elf2-siRNA-transfected cells, while control cells showed recovery of the junction barrier (**Fig. 6f, g**). These findings suggest that Elf2 is critical for maintaining endothelial barrier integrity in HLMVEC.

### EC-restricted deletion of Elf2 (*Elf2^ΔEC^*) in adult mice reduces VE-cadherin and Tie2 expression and exacerbates sepsis-induced lung vascular injury

To investigate the *in vivo* role of Elf2 in EC, we generated EC-restricted Elf2 knockout (*Elf2^ΔEC^*) mice using a Cre-dependent Cas9 knock-in system. CRISPR/Cas9-cdh5-Cre mice were created by crossing Rosa26-LSL-Cas9 knock-in mice with Cdh5-Cre transgenic mice (13,33). *Elf2^ΔEC^* mice were generated through *in vivo* delivery of a plasmid encoding a single guide RNA (sgRNA) targeting the *mElf2* gene, with scrambled sgRNA (Sc-sgRNA) as a control. Administration of the mElf2-sgRNA plasmid effectively reduced Elf2 expression in lung EC (**Fig. 7a, b**). Notably, expression of EC-specific genes VE-cadherin and Tie2 was significantly reduced in *Elf2^ΔEC^* mice compared to controls, while Cd31 and β-catenin levels remained unchanged (**Fig. 7c**). Immunostaining further demonstrated reduced VE-cadherin expression in lung vessels of *Elf2^ΔEC^* mice (**Fig. 7d**). H&E staining of lung sections revealed increased perivascular space in *Elf2^ΔEC^*mice, indicative of basal vascular leakage (**Fig. 7e**). Both basal and LPS-induced vascular leak were significantly enhanced in *Elf2^ΔEC^* mice (**Fig. 7f**), with 100% mortality observed within 24 hours following a low-dose LPS challenge (5 mg/kg, i.p.) in *Elf2^ΔEC^* mice, whereas WT mice showed no mortality (**Fig. 7g**). These findings aligned with previous studies showing that optimal expression of VE-cadherin and Tie2 is essential for a stable endothelial barrier (1–6,14). Based on these results, we propose a model for the epigenetic regulation of endothelial junctional barrier (**Fig. 7h**).

**Fig. 7:**
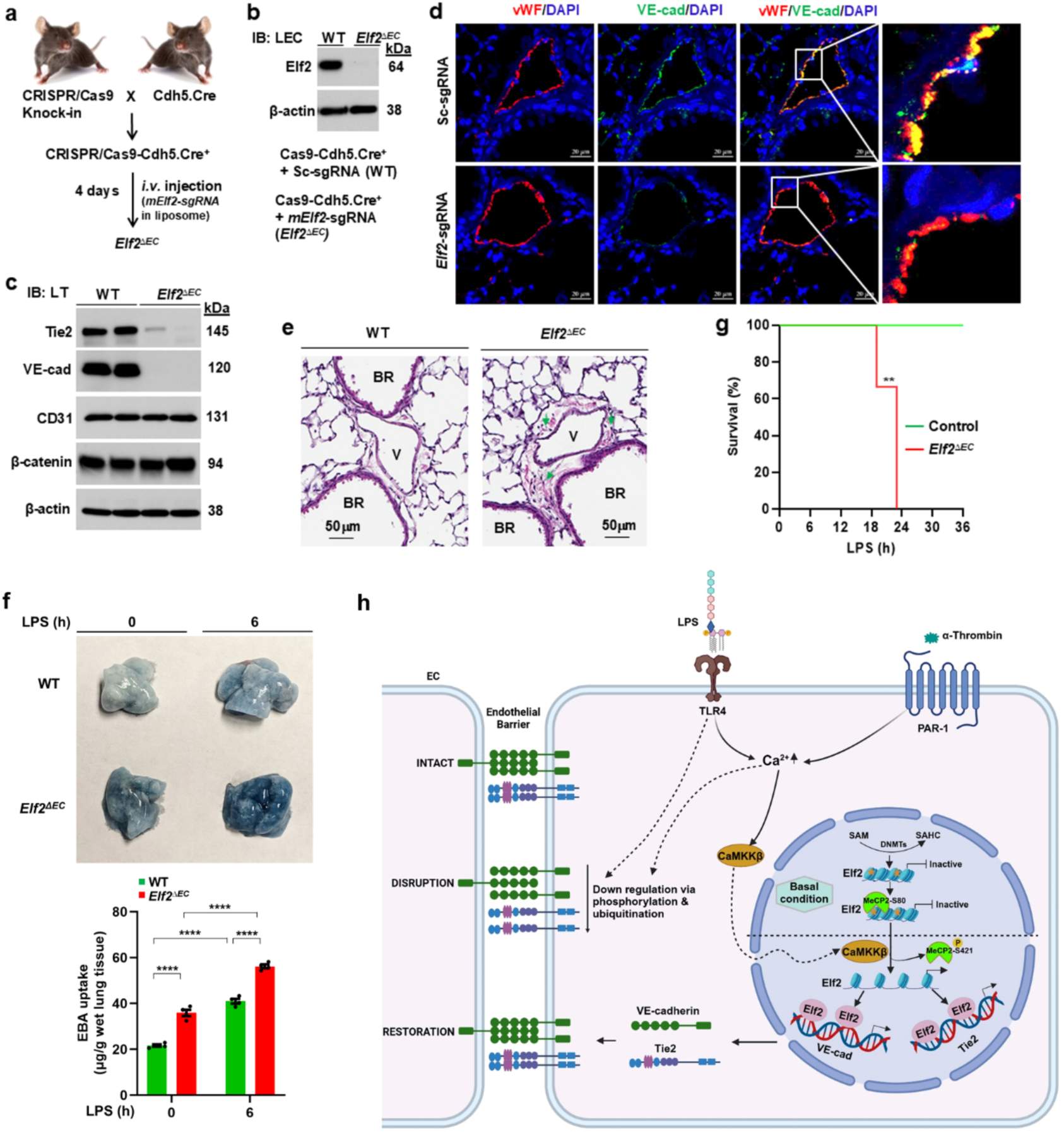
EC-restricted deletion of Elf2 (*Elf2^ΔEC^*) in adult mice reduces VE-cadherin and Tie2 expression and exacerbates sepsis-induced lung vascular injury. a, Depicts the protocol used to create EC-restricted *Elf2* knockout (*Elf2^1EC^*) mice. The mixture of liposome and plasmid expressing sgRNA to target *mElf2* or scrambled sg-RNA (Sc-sgRNA) was injected i.v. into the CRISPR/Cas9-cdh5-Cre mice. Four days after injection, mice were used for experiments. **b**, LEC isolated from mice injected with Sc-sgRNA (WT) or sgRNA to target *mElf2 (Elf2^1EC^*) were used for IB analysis. **c**. IB analysis of lung tissue from WT and *Elf2^1EC^* mice. **d**, lung sections from WT and *Elf2^1EC^*mice were stained with antibodies specific to VE-cadherin, vWF (EC-marker), and DAPI. *Right panels* show the magnified images. **e**. H&E staining of lung sections. BR, bronchi; V, vessel. Green arrow heads showed increased perivascular space in *Elf2^1EC^* mice indicating basal vascular leak. **f**, WT and *Elf2^1EC^* mice were challenged with LPS (5 mg/kg, i.p.) and then used to assess *in vivo* lung vascular leak by measuring EBA uptake. Augmented lung vascular leak was observed in *Elf2^1EC^* mice. N = 4 mice per genotype; ****p< 0.0001 (Two-way ANOVA). **g**, Survival following LPS (5 mg/kg, i.p.) in WT and *Elf2^1EC^* mice. N = 6 in each group. **p< 0.01 (log-rank test). **h**, Model for epigenetic regulation of Elf2 gene expression in endothelial cells during repair of sepsis-induced lung vascular injury created in https://BioRender.com. In quiescent endothelial cells (EC), the CpG-rich promoter of the Elf2 gene is methylated by DNA methyltransferases (DNMTs). This methylation recruits the methyl-CpG binding protein MeCP2, which binds to the methylated CpG sites and represses Elf2 transcription. During endotoxemia, activation of TLR4 and/or PAR-1 increases intracellular Ca²⁺ levels in EC, leading to loss of endothelial barrier integrity through phosphorylation-driven internalization and ubiquitin-mediated degradation of VE-cadherin (5,13). The rise in intracellular Ca²⁺ also activates Ca²⁺/calmodulin-dependent kinase CaMKKβ, which translocates to the nucleus and phosphorylates MeCP2 at Ser^421^. This phosphorylation causes MeCP2 to dissociate from methylated CpG sites, triggering CpG demethylation and reactivation of Elf2 transcription. Once reactivated, Elf2 promotes its own expression and upregulates VE-cadherin and Tie2, facilitating the repair of the disrupted endothelial barrier. SAM, S-adenosyl-methionine; SAHC, S-adenosylhomocysteine; TLR4, toll-like receptor 4; PAR-1, protease-activated receptor 1; Thr, thrombin; orange star, methylated CpGs.

## Discussion

Acute lung injury (ALI) and its most severe manifestation, acute respiratory distress syndrome (ARDS), are leading causes of death in ICU patients. A hallmark of ARDS is the increased lung vascular permeability, resulting in the accumulation of protein-rich edema fluid and inflammatory cells in lung tissue and airspaces (34–37). ALI/ARDS can arise from pneumonia, sepsis, severe trauma, and viral infections such as influenza or SARS-CoV-2 (COVID-19), leading to similar pathological outcomes (34–39). A key feature underlying ALI is the breakdown of the endothelial junctional barrier, where VE-cadherin expression at EC adherens junctions (AJs) is critical for maintaining barrier integrity. The Ang-1/Tie2 signaling axis plays a constitutive role in stabilizing the endothelial barrier by promoting VE-cadherin expression at AJs (6). However, the expression of barrier-stabilizing proteins VE-cadherin and Tie2 is downregulated in inflammatory conditions like bacterial sepsis (13–15), suggesting that the re-expression of these proteins is necessary to restore endothelial barrier integrity following vascular injury.

The Ets family of transcription factors plays a critical role in the transcription of EC-specific genes (17–21). Studies have indicated that the Ets family member Elf2 can bind to and activate the promoters of Tie2 and Cdh5 (VE-cadherin) (20, 21); however, how Elf2 expression in EC is transcriptionally regulated is unclear. In this study, we demonstrate the epigenetic regulation of Elf2 gene transcription through Ca^2+^/calmodulin-dependent kinase CaMKKβ in EC. Using knockout mouse model, genome-wide methylation analysis, gene knockdown studies, and EC-specific knockout mouse models, we show that EC-expressed CaMKKβ prevents methylation of the Elf2 gene, thereby promoting the expression of Elf2 and its target genes, Tie2 and VE-cadherin, to stabilize the endothelial junctional barrier. Our findings reveal the crucial role of CaMKKβ-mediated Elf2 transcription in regulating endothelial AJ integrity.

The Ca^2+/^calmodulin (CaM) complex binds to CaMKKβ, activating its catalytic function. CaMKKβ then activates downstream kinases by phosphorylating them in their activation loops (22,23). CaMKKβ signaling also promotes transcriptional activation through the phosphorylation of transcription factors, e.g., CREB (22,23). Here, we explored the role of EC-expressed CaMKKβ in regulating EC junctional barrier. We found that CaMKKβ-deficient mice (*Camkkβ^−/−^*) were highly susceptible to LPS-induced lung vascular injury and mortality. Consistent with this, VE-cadherin and Tie2 mRNA and protein levels were significantly reduced in *Camkkβ^−/−^* mice, indicating a role for CaMKKβ in regulating endothelial barrier-stabilizing genes.

Since CaMKKβ signaling pathways are known to regulate DNA methylation-mediated repression of gene expression (26), we performed genome-wide methylation analysis of lung tissues from WT and *Camkkβ^−/−^* mice. We found hypermethylation of several genes in *Camkkβ^−/−^* mice, with Elf2 being a top hit. Inhibition of DNA methylation with 5-AZA or expression of WT-CaMKKβ, but not a kinase-defective mutant (K193A-CaMKKβ), restored Elf2, VE-cadherin and Tie2 expression in *Camkkβ^−/−^* mice, supporting the concept that CaMKKβ prevents Elf2 gene methylation. Using EC-specific CaMKKβ knockout mice, we showed that CaMKKβ is required for the expression of Elf2, VE-cadherin and Tie2 in EC. Further, CaMKKβ depletion in HLMVEC using siRNA also suppressed Elf2, VE-cadherin and Tie2 expression, highlighting the essential role of CaMKKβ in regulating these genes. Following LPS challenge in WT mice, we observed a transient reduction in CaMKKβ, Elf2, VE-cadherin and Tie2 expression, which returned to basal levels after 48 hours, suggesting that CaMKKβ expression precedes to promote the re-expression of these genes.

MeCP2 has been shown to induce long-term silencing of gene expression through binding to methyl-CpG enriched promoter regions of genes (26,29). Studies have shown that MeCP2 is highly expressed in EC, and it represses the expression of genes including Kruppel-Like Factor 2 (KLF2) (40), inducible nitric oxide synthase (iNOS) (30), and vascular endothelial growth factor (VEGF) (31). Under basal state, phosphorylation of MeCP2 at S^80^ mediates MeCP2 binding to methylated CpG DNA (26). Increase in intracellular Ca^2+^ causes phosphorylation of MeCP2 at S^421^ to dissociate MeCp2 from methylated CpG DNA and hence promotes gene transcription (26).

We observed that MeCP2 interacts with methylated CpG sites on Elf2 promoter region in HLMVEC under base state. As expected, thrombin stimulation, which increases intracellular Ca^2+^ level, caused dissociation of MeCP2 from the Elf2 gene promoter. Moreover, we observed that either thrombin or LPS challenge caused dephosphorylation of MeCP2 at S^80^ and phosphorylation of MeCP2 at S^421^. Importantly, inhibition of CaMKKβ, blocked thrombin- or LPS-induced effects on MeCP2 in HLMVEC. Further, we observed that in EC-restricted *Mecp2* knockout (*Mecp2^1EC^*) mice, augmented expression of Elf2, VE-cadherin and Tie2. These results together support that MeCP2 represses Elf2 transcription, whereas CaMKKβ derepresses the Elf2 transcription to regulate endothelial barrier integrity.

Although *in vitro* studies have demonstrated that the Ets factor Elf2 can bind and transactivate the promoters of the genes encoding the receptor tyrosine kinase Tie2 and Cdh5 (20,21), the role of EC-expressed Elf2 in regulating endothelial barrier through the expression of VE-cadherin and Tie2 is unknown. In the present study, we first showed that under basal state the binding of Elf2 to the promoter regions of genes encoding Tie2 and Cdh5 in HLMVEC. Next, we demonstrated that depletion of Elf2 in HLMVEC, suppressed the expression of VE-cadherin and Tie2. Basal as well as inflammatory mediators, thrombin- or LPS-induced permeability increase was augmented in Elf2 depleted HLMVEC compared with control HLMVEC. Further, LPS challenge caused loss of VE-cadherin expression in both control and Elf2 depleted HLMVEC; however, in control HLMVEC, VE-cadherin expression was restored to near baseline level within 24 h of LPS challenge, whereas in Elf2 depleted HLMVEC, VE-cadherin expression was not restored, supporting the crucial role of Elf2 in regulating EC barrier. Next, we deleted *Elf2* in adult mouse endothelial cells in which EC-restricted Elf2 knockout (*Elf2^1EC^*) showed defective in the expression of Tie2 and VE-cadherin. Basal as well as LPS-induced lung vascular leak was augmented in *Elf2^1EC^* mice compared with WT type counterparts. In addition, *Elf2^1EC^* mice failed to survive up to 24 h with a low dose of LPS challenge. These results together support the essential role of EC-expressed Elf2 in controlling the endothelial junctional barrier integrity through the expression of VE-cadherin and Tie2 in EC.

In summary, our study demonstrates that CaMKKβ signaling is crucial for maintaining and repairing the endothelial junctional barrier by preventing Elf2 gene methylation and promoting Elf2-mediated expression of VE-cadherin and Tie2. These findings highlight the regulatory roles of CaMKKβ-MeCP2-Elf2 signaling axis in endothelial barrier function, with potential therapeutic implications for conditions involving vascular injury.

## MATERIALS AND METHODS

### Antibodies

Rabbit polyclonal antibody (pAb) against β-catenin (Cat^#^ sc-7199), goat pAb against CaMKKβ (Cat^#^ sc-9629), mouse mAb against Elf2/Nerf2 (Cat^#^ sc-130632), mouse mAb against VE-cadherin/cdh5 (Cat^#^ sc-9989), and goat pAb against vWF (Cat^#^ sc-8068) were from Santa Cruz Biotechnology (Santa Cruz, CA). Rabbit pAb against CaMKKβ (Cat^#^ ab96531), rabbit pAb against Elf2/Nerf2 (Cat^#^ ab225958), rabbit pAb against MeCP2 (Cat^#^ ab2828), and rabbit pAb against VE-cadherin (Cat^#^ ab33168) were from Abcam (Cambridge, MA). Mouse mAb against β-actin (Cat^#^ A5441), rat mAb against PECAM-1 (CD31) (Cat^#^ CBL1337), mouse mAb against Tie2 (Cat^#^ 05-584), and rabbit mAb against vWF (Cat^#^ AB7356) were from Millipore Sigma/EMD Millipore Corp. (St. Louis, MO). HRP-conjugated mouse mAb against β-actin (Cat^#^ HRP-66009), and rabbit pAb against Lamin B1 (Cat^#^ 12987-1-AP) were from Proteintech Group (Rosemont, IL). Rabbit pAb against phospho-MeCP2 (Ser80) (Cat^#^ PIPA513024, Cat^#^ PIPA5143716), rabbit pAb against phospho-MeCP2 (Ser421) (Cat^#^ PA5-35396), Dynabeads Sheep anti-rat IgG (Cat^#^ 11035), donkey anti-goat horseradish peroxidase conjugated secondary antibody (Cat^#^ A16005), Alexa Fluor 594 chicken anti-rabbit secondary antibody (Cat^#^ A21442), Alexa Fluor 488 goat anti-mouse secondary antibody (Cat^#^ A11001), Alexa Fluor 647 chicken anti-mouse secondary antibody (Cat^#^ A21463), Alex Fluor 488 chicken anti-rabbit secondary antibody (Cat^#^ A21441), and Alexa Fluor 488 goat anti-rabbit secondary antibody (Cat^#^ A11034) were from Invitrogen by Life Technologies (Carlsbad, CA). KPL peroxidase-labeled anti-mouse secondary antibody (Cat^#^ 5450-0011) and KPL peroxidase-labeled anti-rabbit secondary antibody (Cat^#^ 5220-0458) were from SeraCare (Milford, MA).

### Other reagents

DNA methyltransferase inhibitor 5-azacitidine (Cat^#^ PHR1911) and Collagenase from *Clostridium histolyticum* (Cat^#^ C2674) were from EMD Millipore. CaMKKβ inhibitor 7-Oxo-7H-benzimidazo [2,1-a]benz[de]isoquinoline-3-carboxylic acid acetate (STO-609) (Cat^#^ 1551) was from Tocris Bioscience, part of Bio-Techne (Minneapolis, MN). PAR-1-activating peptide (TFLLRNPNDK-NH2) was custom synthesized as the C-terminal amide with a purity of 97.2% by Genscript (Piscataway, NJ). CaMKKβ WT plasmid (Cat^#^ 33322) and CaMKKβ K193A mutant plasmid (Cat^#^ 33323) were from Addgene (Watertown, MA). AllStars negative control siRNA (Sc-siRNA) (Cat^#^ 1027281) was from Qiagen (Hilden, Germany). SiRNA transfection reagent (Cat^#^ sc-29528), human (*h*)-specific siRNA to target CaMKKβ (Cat^#^ sc-38955), and Elf2/Nerf2 (Cat^#^ sc-43961) were from Santa Cruz Biotechnology. Lipopolysaccharides (LPS) from *Escherichia coli* O111:B4 (Cat^#^ L3012), Chloroform (Cat^#^ 288306), Cholesterol (Cat^#^ C8667), Dimethyldioctadecylammonium bromide (Cat^#^ D2779), Evans Blue (Cat^#^ E2129), Dimethyl Sulfoxide (DMSO, Cat^#^ D2650), and Sodium citrate (Cat^#^ S-4641) were from Sigma-Aldrich Inc (St. Louis, MO). Human α-thrombin (Cat^#^ HT1002A) was from Enzyme Research Laboratories Inc. (South Bend, IN). Fetal bovine serum (FBS) (Cat^#^ 33323) was from HyClone Laboratories, Cytiva (Marlborough, MA). 0.05% Trypsin-EDTA solution (REF^#^ 25300-054) and Penicillin/Streptomycin in 10,000 U/mL (Cat^#^ 15140-122) were from Gibco, by Life Technologies. Triton X-100 (Cat^#^ BP151) and Tween 20 (Cat^#^ BP337) were from Fisher Scientific (Hampton, NH). Gelatin (Cat^#^ 170-6537) was from BIO-RAD Laboratories (Des Plaines, IL). Horse serum (Cat^#^ 100-508) was from Gemini Bio (West Sacramento, CA).

### Endothelial cells (EC)

HLMVEC obtained from Lonza Walkersville Inc. (Cat^#^ CC-2527; Walkersville, MD) or from Cell Biologics (Cat^#^ H-6011; Chicago, IL) were used in this study. EC were grown in EBM medium (Cat^#^ CC-3156) supplemented with EGM^TM^-2 MV Microvascular Endothelial SingleQuots^TM^ Kit (Cat^#^ CC-4147), 10% FBS and penicillin/streptomycin solution at 37°C incubator with 5% humidified atmosphere of CO_2_. The EC from both sources were used between passages 4 and 6 for experiments. Mouse EC was isolated from lung tissues as described by us (41) and used directly for experiments without further culture.

### Mice

C57BL6/J mouse strain (Code^#^ 027) was from Charles River Laboratories (Wilmington, MA). With C57BL/6 background, VE-cadherin-Cre (cdh5-Cre) mouse strain B6.FVB-Tg(Cdh5-cre)7Mlia/J (Stock 006137), CaMKKβ^+/−^ (heterozygous) mouse strain B6.129X1-*Camkk2^tm1Tch^*/J (Stock 014172), Mecp2 floxed (*Mecp2^fl/^*^fl^) mouse strain B6;129P2-Mecp2^tm1Bird^/J (Stock 006847), and CRISPR/Cas9 knock-in mouse strain Rosa26-LSL-Cas9 knock-in on B6j (Rosa26-floxed STOP-Cas9 knock-in on B6J) (Stock 026175) were from The Jackson Laboratory (Bar Harbor, ME). *Mecp2*^fl/fl^ mice were bred with Cdh5-Cre mice to generate *Mecp2^fl/fl^-Cre^-^* (WT), *Mecp2^fl/+^-Cre^-^*(WT) and Mecp2^fl/fl^-Cre^+^ (*MeCp2^1EC^*) mice. CRISPR/Cas9 knock-in mice were bred with Cdh5-Cre mice to obtain Cas9 and Cdh5-Cre double positive progenies. *Elf2^ΔEC^*mice were generated via CRISPR/ Cas9 gRNA system as described previously (13). The *m*Elf2 CRISPR sgRNA (5′-CGACGAGACTTATATGATGC-3′) and non-targeting control sgRNA (5′-GCGAGGTATTCGGCTCCGCG-3′) in pGS-gRNA vector were custom prepared by GenScript. Camkkβ expression in mouse lung EC was disrupted by liposome-mediated delivery of CRISPR/Cas9 system (pSPCas9 BB-2A-GFP PF458 (42). This CRISPR plasmid expresses Cas9 under the control of the mouse Cdh5 promoter (EC-specific) and sgRNA driven by the U6 promoter. This plasmid encodes control-sgRNA (5′-GCGAGGTATTCGGCTCCGCG-3′) or *m*Camkkβ-sgRNA (5′-AAACGTCGGATCAGCTTCTTTTTGC-3′). Liposome was prepared as described previously (43) by mixing Cholesterol and Dimethyldioctadecylammonium bromide (1:1 molar ratio) in Chloroform and dried using the Buchi Rotavapor R-300 (New Castle, DE). The dried lipid was dissolved in 5% dextrose followed by 30-min sonication in a bath sonicator Branson 2510 Ultrasonic (Danbury, CT). The complex consisting of plasmid DNA and liposomes were combined at a ratio of 1 μg of DNA to 8 nmol of liposomes. A total volume of 150 µl liposome/plasmid complex containing 30 µg of sgRNA plasmid was retro-orbitally injected into each WT or Cas9-Cdh5-Cre^+^ mice to generate experimental control or EC-specific knockout mice. Four 4 days post injection, mice were used for experiments. 5-Aza at the dose of 1 mg/kg was intraperitoneally injected to CaMKKβ^+/+^ (WT) and *CaMKKβ^-/-^* mice once per day for 5 consecutive days. All mice were maintained in pathogen-free environment at the University of Illinois Animal Care Facility in accordance with institutional guidelines of the National Institutes of Health (NIH). All animal experiments were performed under the protocol approved by the Institutional Animal Care and Use Committee of the University of Illinois at Chicago.

### Lung injury in mice

Eight to twelve weeks old both male and female mice were intraperitoneally injected (i.p.) with a single dose of LPS at 5 mg per kg body weight otherwise indicated. *In vivo* lung injury was assessed by measuring Evans blue dye bond albumin (EBA) uptake in lungs (13, 44), H&E staining, and determining lung myeloperoxidase (MPO) activity. For histology, paraffin-embedded sections in 5 μm thickness were prepared from the lungs by the Research Histology Core at UIC Research Resources Center (RRC). For MPO assay, lung tissues were frozen in liquid nitrogen directly after their removal from mice. Following hexadecyltrimethylammonium bromide (HTAB) extraction and dianisidine-H_2_O_2_ reaction (45), the MPO levels were determined based on the absorbance changes at 450 nm over a 3-min period in Beckman DU530 spectrophotometer (Fullerton, CA). Polymicrobial sepsis was induced by cecal ligation and puncture (CLP) as described by Rittirsch et al. (46). The caecum was punctured using 18-gauge needle on five different places. For survival studies, mice were monitored four times daily.

### CpG island regions and CpG sites prediction

The promoter sequence for mouse and human genes was obtained from the EPDnew databases (https://epd.expasy.org/epd/EPDnew_database.php). Then, the web-based program DBCAT (Database of CpG Islands and Analytical Tools) was used to map CpG island regions and CpG sites in these promoter regions. The analysis parameters were set as follows: Observed to Expected ratio (o/e) as 0.60, minimal length as 200, any CpG island with GC content less than 50% filtered out.

### Methylated DNA Immunoprecipitation (MeDIP)-sequencing and data analysis

Basal lung tissue from WT and CaMKKβ^-/-^ mice were used for genomic DNA isolation using the Qubit dsDNA HS reagent (Invitrogen), followed by a purification using the Genomic DNA Clean & Concentrate kit (Zymo Research). The purified genomic DNA was sheared and used as input for the Methylated DNA IP kit (Zymo Research), along with a spike-in Control DNA. Samples were processed as per the protocol. IP DNA as well as input control DNA for each sample were processed in parallel for library construction and sequencing using the Pico Methyl-Seq Library Prep kit (Cat^#^ D5455) by following the manufacturer’s instructions (Zymo Research). Libraries were combined into 2 pools and analyzed on a TapeStation HS D1000 tape (Agilent). Sequencing was carried out on NextSeq 500 (Illumina), 2×42 nt reads, High Output Kit (400 million reads).

The raw data were aligned to the mm10 reference genome using BWA mem (47). Apparent PCR duplicate reads were removed from further analysis using picard tools (http://broadinstitute.github.io/picard/). Peaks for each IP sample were called against its matching non-pulldown control (input) using MACS2 (48) with default parameters. Peaks with a score >5 (p-value <1e^-5^) were retained. The peaks in promoters (+/-2kb of the TSS) were filtered from further analysis. Prior to differential analysis, peak calls across all samples were merged using bed tools merge (49) and read counts per peak were quantified for both IP and input samples using feature counts (50,51). Input read counts were subtracted from their paired IP count after adjusting for differences in sequencing depth by stochastically up- or down-sampling the input counts according to a binomial model. EdgeR was used to perform differential analysis between two sample groups to identify the genes with significantly different Methylation levels in TSS regions (52,53).

### Promoter analysis and ChIP assay

The binding sites for transcription factors in 5′-regulatory regions of genes were located via either Genomatix (München, Germany), Eukaryotic promoter Database (SIB; https://epd.epfl.ch//index.php) or via searching binding motif sequences manually. ChIP assays were performed by using ChIP assay kit (Cat^#^ 17-295) from Millipore. The primers used for successfully detecting binding regions in specific genes were custom synthesized by IDT and listed as follows: Elf2 gene binding on Tie2 promoter (region −309 – −68: F 5’-AACCAACAGGCCATCTGTGT, R 3’-GGGGCTACTGGGATCTCTGA; region −70 – +171: F 5’-CCCGGGAGAGCTGTTAGAAG, R 3’-CAAGCCCTATCCATCTCGCA; region +42 − +281: F 5’-TTCTGAAAATGCTGACCGGG, R 3’-CTGTCTGAGCACAGGGAGTTT; region +363 – +576: F 5’-TGGAAAGTCACAAACCGCTG, R 3’-TGGGGAGAGCACCCAGAAAT; region +561 – +685: F 5’-CTGGGTGCTCTCCCCAAATC, R 3’-TGCCACAGAGCCTTTGCATT). Elf2 gene binding on VE-cadherin promoter (region +239 – +332: F 5’-CACTCCCATGACAGAGAGGC, R 3’-CTTGGCACTACCTCTGGGTG; region +575 – +648: F 5’-CCCCTACCCATACGCCACTA; R 3’-CAAGGCTCGGAGCTTCCTTT; region +835 – +1034: F 5’-GGCTAGAGCATAAGAGCCCC, R3’-TAATGAGCCCCACACCATCC). MeCP2 binding on Elf2 promoter (region −562 − −308: F 5′-TTGGGGATGCGGTGACG-3′, R 5′-TTCTCCCAAACCCGCCT-3′).

### Quantitative real-time PCR (QRT-PCR)

Total RNA was extracted from mouse lung tissues by using RNeasy Mini Kit (Cat^#^ 74104) from Qiagen or TRIzol Reagent (REF^#^ 15596026) from Ambion by Life Technologies. The complementary DNAs was reverse transcribed with Oligo (dT)_18_ from these RNA samples by using RevertAid First Strand cDNA Synthesis Kit (Cat^#^ K1621) from Thermo Fisher Scientific (Waltham, MA). The primers used for detecting gene expression in these samples were custom synthesized by Integrated DNA Technologies (IDT) (Coralville, IA): Angiopoietin (Ang)-1 [forward (F) 5′-TTGTGATTCTGGTGATTGTGG-3′, reverse (R) 5′-CTTGTTTCGCTTTATTTTTGT-3′)], Ang-2 (F 5′-GACTTCCAGAGGACGTGGAAAG-3′, R 5′-CTCATTGCCCAGCCAGTACTC-3′), β-catenin (F 5′-ACTGCTGGGACTCTG-3′, R 5′-TGATGGCGTAGAACAG-3′), CaMKKβ (F 5′-CAGCTCTGTGGGTTCGAGTT-3′, R 5′-GGGAGTCAAACAGAAGCGGA-3′), Elf1 (F 5′-TGTCCAACAGAACGACCTAGT-3′, R 5′-CACACAAGCTAGACCAGCATAA-3′), Elf2 (F 5′-TTTGTCTCGTCCTGTGGTGC-3′, R 5′-TACTGCTCTCCCTCCACGTC-3′), Occludin (F 5′-ACTGGGTCAGGGAATATCCA-3′, R 5′-TCAGCAGCAGCCATGTACTC-3′), VE-cadherin (F 5′-ACGGACAAGATCAGCTCCTC-3′, R 5′-TCTCTTCATCGATGTGCATT-3′), VEGFR-1 (F 5′-GAGGAGGATGAGGGTGTCTATAGGT-3′, R 5′-GTGATCAGCTCCAGGTTTGACTT-3′), VEGFR-2 (F 5′-GCCCTGCTGTGGTCTCACTAC-3′, R5′-CAAAGCATTGCCCATTCGAT-3′), Tie2 (F 5′-AAGCATGCCCATCTGGTTAC-3′, R 5′-GCCTGCCTTCTTTCTCACAC-3′), β-actin (F 5’-ACTCCTATGTGGGTGACGAG-3’, R 5’-ATCTTTTCACGGTTGGCCTTAG-3’), GAPDH (F 5’-ACCCAGAAGACTGTGGATGG-3′, R 5′-CACATTGGGGGTAGGAACAC-3′). The quantitative real-time PCR reactions were prepared by using Fast SYBR Green Master Mix (Cat^#^ 4385612), and the amplification was performed in Quant studio7 Flex Real-Time PCR System (Applied Biosystems). The results were calculated using the comparative CT (ΔΔCT) formula and normalized with the expression of housekeeping genes β-actin or GAPDH.

### siRNA transfection

EC grown to 70–80% confluence on gelatin-coated culture dishes were transfected with gene specific siRNAs or Sc-siRNA by following Santa Cruz’s instructions. At 72 h post transfection, cells were used for experiments.

### Transendothelial monolayer electrical resistance (TER) measurement

As described in Tiruppathi et al. (54), the real-time changes in transendothelial monolayer electrical resistance (TER) were measured to assess endothelial barrier function in ECIS1600R from Applied Biophysics (Troy, NY). HLMVEC around 80% confluence were transfected either with Sc-siRNA or Elf2-siRNA. At 24 h post transfection, the cells were detached from the culture dish by using Trypsin and seeded on the ECIS 8W1E PET from Applied Biophysics. The confluent EC monolayer incubated in basal medium with 2% FBS was exposed to indicated PAR-1 peptide or LPS. Data are presented as resistance normalized to its stable starting value.

### IB analysis

The confluent EC were treated with or without specific agents and lysed in RIPA lysis buffer (50 mM Tris-HCl, pH 7.5, 150 mM NaCl, 1 mM EGTA, 1% Triton X-100, 0.25% sodium deoxycholate, 0.1% SDS, 10 μM orthovanadate, and protease-inhibitor mixture). Mouse lungs were homogenized in the lysis buffer as used in EC. The lysate from either EC or mouse lungs were centrifuged at 13,000 rpm for 10 min to collect supernatant for IB analysis. The total proteins for either EC lysates or lung lysates were resolved on a 4-15% gradient SDS-PAGE gel and transferred to a polyvinylidene difluoride (PVDF) membrane. The membrane was blocked with 5% dry milk in TBST (10 mM Tris-HCl, pH 7.5, 150 nM NaCl, and 0.2% Tween-20) at RT for 1 h, and then probed with the indicated primary antibody diluted with 2.5% milk or 1% BSA in TBST overnight at 4°C. Next day, the membrane was washed three times with TBST and then incubated with appropriate peroxidase-labeled secondary antibody. Protein bands were detected by enhanced chemiluminescence and the target IB bands were quantified using NIH ImageJ software.

### Immunofluorescence (IF) staining

EC grown on the 0.2% Gelatin coated glass coverslips were fixed with 2% paraformaldehyde for 10 min and permeabilized with 0.05% Triton X-100 for 5 min at 4°C. Then, the cells were blocked in PBS containing 5% horse serum and 1% BSA for 1 h at room temperature (RT). After blocking, the cells were incubated with specific primary antibody overnight at 4°C. Next day, the cells were incubated with Alexa-Fluor-conjugated secondary antibodies for 1 h at RT. Finally, the cells were mounted with ProLong (R) Gold antifade with 4′, 6-diamidino-2-phenylindole (DAPI) (Cat^#^ 8961S) from Cell Signaling Technology (Danvers, MA). In each step, cells were washed three times with PBS.

The paraffin-embedded lung tissue sections were heated on a platform at 90°C until melted, followed by the dewax step in xylene and rehydration step in a series of 100% ethanol, 95% ethanol, 75% ethanol, 50% ethanol and water. Then, the sections were boiled in sodium citrate buffer (10 mM citrate, 0.05% Tween 20, pH 6.0) for 25 min to unmask antigens. At RT, the sections were permeabilized in 0.3% Triton X-100 in PBS for 10 min. After blocking with 5% horse serum, 1% BSA and 0.2% Triton X-100 in PBS for 1 h, the sections were incubated in indicated primary antibodies in PBS containing 1% BSA at 4°C for overnight. Next day, the sections were re-probed with corresponding fluorochrome coupled secondary antibodies at RT for 1 h. In the last step, the sections were mounted in ProLong gold antifade coupled with DAPI. Images were acquired with the Zeiss LSM 880 confocal microscope.

### Statistical analysis

Statistical analyses were performed either in GraphPad Prism (Boston, MA) or in Excel. ANOVA, unpaired Student’s t test (two-tailed), and log-rank test were used to determine statistical significance. The *p* value < 0.05 was considered significant.

## Acknowledgments

This work was funded by NIH grants R01HL156965 and P01HL160469-project 2. M.M.C. at Research Informatics Core supported in part by NCATS grant UL1TR002003. The schematic diagrams were made using Biorender.com.

## Author contributions

D.M.W.: conceived, created multiple mouse models, study design and execution, data analysis and interpretation, and manuscript writing; J.Y.: experimental work; M.O.A.: experimental work; Z.A.: performed MeDIP-sequencing; M.M.C.: performed MeDIP-sequencing data analysis; V.N. & A.B.M.: fund acquisition and study design; C.T.: fund acquisition, conceived, study design, data interpretation, supervision, and manuscript writing.

