## Supplemental Data Table 1 for "CaMKKβ regulates transcription factor Elf2 gene methylation to maintain endothelial junctional barrier integrity"

**MeDIP-seq analysis of lung tissue from WT and KO (*Camkkβ*$^{-/-}$) mice**

| Peak | WT/KO : logFC | WT/KO : logCPM | WT/KO : pvalue | WT/KO : Qvalue | Chr | 2kb up of TSS | 2kb down of TSS | ensembl gene |
|---|---|---|---|---|---|---|---|---|
| peak_8975 | -10.10850694 | 6.0987497 | 1.71E-17 | 2.98E-13 | chr14 | 65803836 | 65807836 | Pbk |
| peak_17573 | -4.179907518 | 5.81348219 | 5.80E-10 | 5.05E-06 | chr3 | 51338658 | 51342658 | Elf2 |
| peak_12185 | 2.484678724 | 10.62894006 | 2.21E-08 | 0.000128297 | chr17 | 39843073 | 39850827 | Gm26917 |
| peak_1761 | 2.360984767 | 6.957086311 | 2.78E-07 | 0.001210518 | chr1 | 195239007 | 195243007 | Gm27940 |
| peak_10801 | 2.117916253 | 6.470727641 | 6.56E-06 | 0.022840656 | chr16 | 35981230 | 35985230 | Gm15564 |
| peak_5478 | -8.431382113 | 4.50258293 | 1.58E-05 | 0.045708387 | chr11 | 120482612 | 120494394 | Mrpl12 |
| peak_19781 | -2.226712881 | 6.510230813 | 3.78E-05 | 0.093945443 | chr4 | 89214863 | 89218863 | ENSMUSG00000083881 |
| peak_26035 | -2.109706567 | 6.637963949 | 4.94E-05 | 0.107429133 | chr7 | 59505276 | 59621095 | Gm24495 |
| peak_951 | 1.976579965 | 6.066163111 | 6.82E-05 | 0.131826611 | chr1 | 113716047 | 113720047 | Gm24937 |
| peak_20904 | 3.171692957 | 4.802357701 | 0.000115276 | 0.200648648 | chr4 | 145444739 | 145448739 | Gm13230 |
| peak_29474 | 8.461955506 | 3.806911564 | 0.000162671 | 0.257405041 | chr9 | 56221771 | 56225771 | Gm24270 |
| peak_31045 | -8.077202638 | 4.246605459 | 0.000306442 | 0.444494235 | chrX | 52910265 | 52914265 | Phf6 |
| peak_15189 | -1.717122539 | 7.976504903 | 0.00034564 | 0.462785915 | chr2 | 69870202 | 69874202 | Gm13625 |
| peak_9579 | -8.108678305 | 4.204844814 | 0.00039333 | 0.489021496 | chr15 | 52476228 | 52480228 | Gm10020 |
| peak_8714 | -2.131468714 | 5.667828801 | 0.00044344 | 0.514567232 | chr14 | 53467755 | 53471755 | Trav8-1 |
| peak_20478 | -4.301123297 | 4.573870467 | 0.00049279 | 0.536093516 | chr4 | 129456350 | 129463580 | Gm24621 |
| peak_29660 | -8.009461244 | 4.097794309 | 0.000935977 | 0.958330248 | chr9 | 69449035 | 69458943 | Gm4978 |
| peak_14918 | -3.023988592 | 4.805713414 | 0.001013893 | 0.980434353 | chr2 | 38971946 | 38975946 | Gm13474 |

Top 18 significantly methylated genes in *Camkkβ*$^{-/-}$ mice were listed.
